# Epithelial HO-1 regulates iron availability and promotes colonic tumorigenesis in a context-dependent manner

**DOI:** 10.1101/2024.03.06.583112

**Authors:** Rosemary C. Callahan, Jillian C. Curry, Geetha Bhagavatula, Alyse W. Staley, Rachel E. M. Schaefer, Faiz Minhajuddin, Liheng Zhou, Rane M. Neuhart, Shaikh M. Atif, David J. Orlicky, Ian M. Cartwright, Mark E. Gerich, Calen A. Steiner, Arianne L. Theiss, Caroline H. T. Hall, Sean P. Colgan, Joseph C. Onyiah

## Abstract

Induction of heme oxygenase-1 (HO-1/*Hmox1*) is broadly considered cytoprotective, but the role of colonic epithelial HO-1 in colitis-associated tumorigenesis is poorly defined. HO-1 catabolizes heme, releasing ferrous iron, a key driver of oxidative stress and lipid peroxidation. We observed that colonic epithelial HO-1 is induced during colitis and tumorigenesis. We also found that HO-1 is upregulated in ferroptosis-inducing conditions in murine and human colonic epithelial organoids, and correlated with lipid peroxidation and ferroptosis markers in colonic tumors. In colonic epithelial organoids exposed to heme, deletion of *Hmox1* amplified a compensatory oxidative stress and detoxification transcriptional program, likely reflecting unresolved oxidative and non-oxidative toxicity from heme. In vivo, epithelial HO-1 deficient mice developed significantly fewer and smaller tumors compared to littermate controls in a colitis-associated tumorigenesis model, despite similar inflammatory injury. Tumors from knockout mice exhibited reduced iron levels, decreased lipid peroxidation, lower oxidative DNA damage, and decreased proliferation. Single-cell RNA sequencing of tumor epithelial cells revealed a shift from a proliferative to a stress-adaptive program with loss of HO-1. These findings identify epithelial HO-1 as a context-dependent regulator of tumorigenesis: protective against acute heme toxicity, but promoting iron-dependent oxidative damage and proliferation in the setting of chronic inflammation.

## Introduction

Inflammatory bowel disease (IBD) is characterized by chronic, relapsing-remitting mucosal inflammation that drives cycles of epithelial injury and repair, significantly increasing the risk of dysplasia and colorectal cancer (CRC) (1-3). This risk is particularly elevated in patients with ulcerative colitis (UC), where disease activity, extent, and duration correlate with neoplastic transformation (4-6). A hallmark of active colitis is mucosal hemorrhage, which exposes the intestinal epithelium to elevated levels of luminal heme released from damaged extravascular erythrocytes (7, 8). Heme is a potent pro-oxidant molecule capable of generating reactive oxygen species (ROS) via its redox active iron atom, leading to oxidative DNA damage, lipid peroxidation, and cell death (9-12).

To mitigate heme toxicity, tissues rely on binding proteins and enzymatic degradation pathways (13). Heme oxygenase-1 (HO-1/Hmox1) is a heme and stress-inducible enzyme that catabolizes heme into biliverdin, carbon monoxide, and ferrous iron (14). Systemic deficiency of HO-1 in humans and mice results in increased oxidative stress, inflammation and tissue injury (15-17). While HO-1 is broadly considered cytoprotective and anti-inflammatory, its role in cancer is complex and context-dependent (14). In the acutely inflamed colon, HO-1 is upregulated in response to inflammation and hemorrhage, but the consequences of its chronic activation, particularly in epithelial cells, have not been well defined. In particular, the intersection of HO-1 activity with iron-regulated oxidative stress and transcriptional responses in the context of colitis-associated cancer (CAC) remains unexplored.

Iron-dependent lipid peroxidation and oxidative stress have been increasingly implicated in both inflammatory and neoplastic processes in the gut, with ferroptosis representing one potential outcome of dysregulated iron metabolism (18-21). Iron metabolism is increasingly recognized as a critical determinant of tumor behavior, with labile iron shown to support nucleotide synthesis, metabolic reprogramming, and proliferation (22, 23). Catabolism of heme by HO-1, with the release of ferrous iron, may therefore have an unintended consequence: fueling tumor-promoting pathways during chronic inflammation. Conversely, loss of HO-1 may exacerbate acute heme toxicity through oxidative and non-oxidative damage from uncatabolized heme, independent of iron release (24). This paradox underscores the need to understand the influence of epithelial HO-1 on colitis and tumorigenesis.

To address this, we utilized a murine model of colitis-associated cancer (CAC) and generated intestinal epithelial-specific HO-1 knockout mice (*Hmox1*^ΔIEC^). We examined the impact of HO-1 deletion on tumor burden, oxidative stress, and compensatory transcriptional responses using epithelial organoids, tissue-level assays, and single-cell transcriptomics. Our findings reveal a complex role for epithelial HO-1 in colitis-associated tumorigenesis: while it protects against acute heme stress, its activity may promote oxidative damage and epithelial proliferation in chronic inflammation through increased iron availability. In its absence, heme triggers a compensatory stress-adaptive transcriptional program that constrains tumor growth.

## Results

### Loss of colonic epithelial HO-1 amplifies stress-linked transcriptional responses to heme

We examined mucosal bleeding as a feature of active colitis using the murine dextran sodium sulfate-induced (DSS) colitis model. We observed a temporal increase in disease activity, fecal bleeding, and fecal heme content, which declined after DSS withdrawal (Fig. 1A-C) (25). Induction of the heme-inducible HO-1/CO pathway has been shown to be protective in murine colitis (26-28). Colonic *Hmox1* mRNA and HO-1 in CD45^-^EpCAM^+^ IECs increased during the course of DSS injury, correlating with bleeding severity and fecal heme (Fig, 1D-E). To investigate colonic epithelial responses to heme we generated mice with IEC specific deletion of *Hmox1* expression (*Hmox1*^ΔIEC^ or knockout/KO mice; Fig. S1A-B) and derived colonic epithelial stem cell organoids (colonoids) from control (*Hmox1*^fl/fl^) and knockout animals. Colonic epithelial cells are relatively resistant to hemin-induced cytotoxicity but upon exposure to moderate amounts of heme we found that loss of *Hmox1* led to increased cell death in murine colonoids (Fig. 1F), similar to a prior report using human colonic epithelial cell lines (10). RNA sequencing revealed distinct transcriptional responses to heme in KO versus control colonoids, with ∼800 genes upregulated in KO and ∼900 in controls (Fig. 1G-I). Kyoto Encyclopedia of Genes and Genomes (29) pathway analysis demonstrated enrichment of glutathione metabolism, ferroptosis-response and several detoxification pathways in KO colonoids (Fig. 1J-L), consistent with increased activation of a broad stress response. Similar to glutathione metabolism, many of the upregulated ferroptosis-response genes are involved in the protective response against iron-induced oxidative damage, such as *Fth1* and *Ftl1*, which store ferrous iron inert as ferritin, or *Slc7a11* and *Slc3a2* that together encode the cystine transporter that is key to maintaining antioxidative glutathione levels.

**Figure 1.**
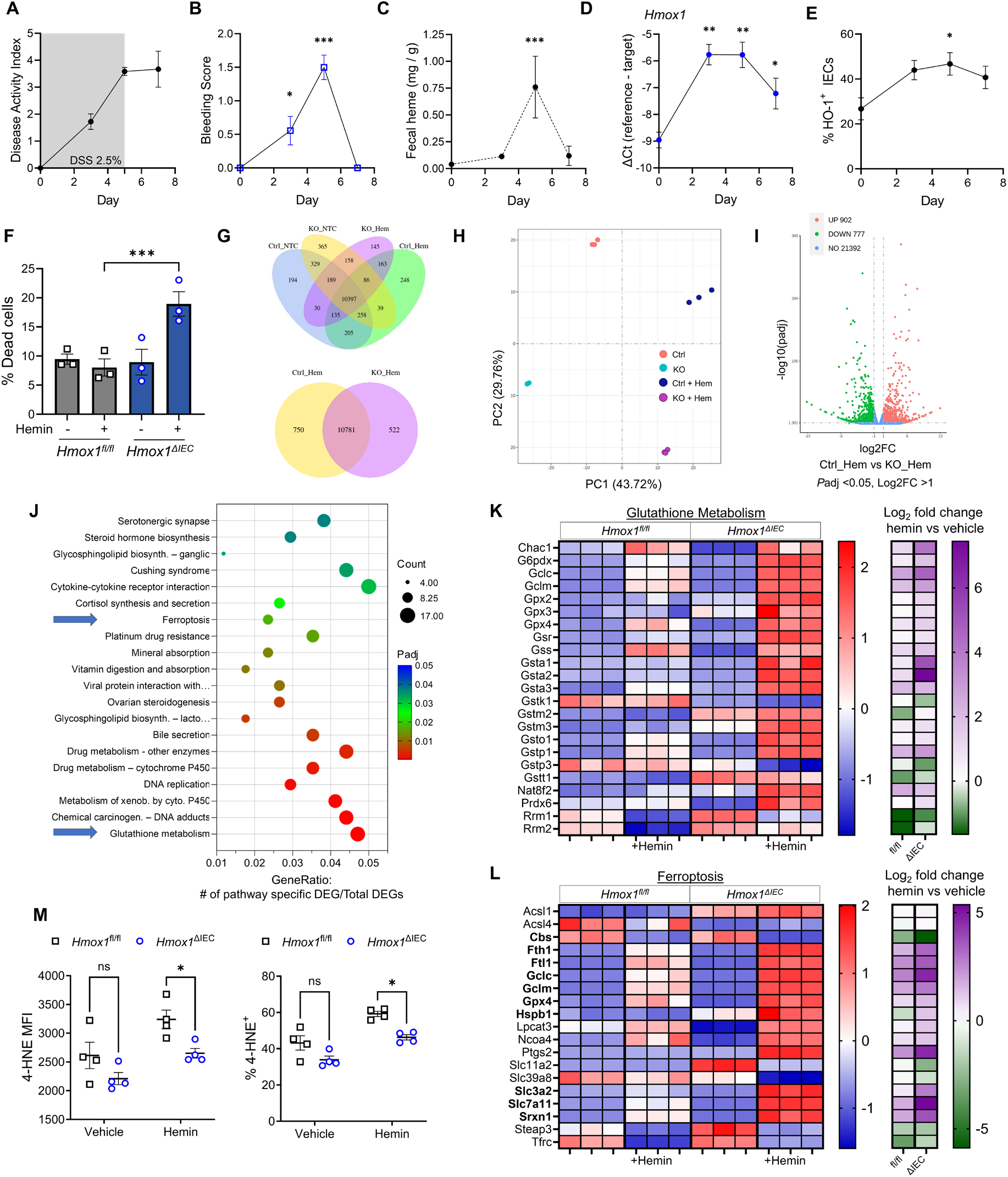
Influence of HO-1 on heme-linked stress responses in colonic epithelial cells. WT mice were given DSS 2.5% in their drinking water for 5 days with daily monitoring up to 2 days after DSS was removed (n = 3-9 per time point). (A) Disease Activity Index (DAI) scoring (weight, stool blood and diarrhea). (B) Bleeding component of DAI score. (C) Fecal heme quantification during course of DSS injury (n = 4-12 per group). (D) Whole colon mRNA expression of *Hmox1* by RT-qPCR (n = 3-10 per group). (E) HO-1 detection using intracellular flow cytometry; expressed as a percentage of total IECs (n = 3 per group). (F) After exposure to hemin (200 µM) for 24h, colonoid cell death was assessed by flow cytometry detection of 7-AAD. (G) Venn diagrams of differentially expressed gene (DEG) patterns in KO and control colonoids in response to hemin, as measured by RNA sequencing. (H) Principal component analysis 2D plot of colonoid groups. (I) Volcano plot of differentially expressed genes (Up = increased expression in control colonoids). (J) KEGG enrichment analysis showing the top 20 significant upregulated pathways in the KO colonoids treated with hemin (Benjamini and Hochberg FDR correction). (K-L) Z-score of DEGs from RNA seq of control and KO colonoids. Log2 fold change of comparison between vehicle and hemin treated groups, by cell type. (L) Bolded genes are typically protective against ferroptosis. (M) Colonoids were exposed to 100 µM of hemin for 24h, and median fluorescence intensity (MFI) and percent 4-HNE positive cells were determined using flow cytometry. Data represent mean ± SEM; \**P* < 0.05, \*\**P* < 0.01, \*\*\**P* < 0.001, and \*\*\*\**P* < 0.0001, by unpaired, Student’s *t* test or one-way ANOVA for multiple comparisons (e.g., compared to day 0 or normal colon).

qPCR analysis confirmed upregulation of antioxidant and ferroptosis-response genes in KO colonoids (Fig. S1C). Depletion of glutathione and inhibition of HO-1 triggers cell cycle arrest in epithelial cells (10, 30, 31). We performed cell cycle analysis and observed increased G0/G1 arrest in KO colonoids exposed to heme (Fig. S1D). These findings suggest that HO-1 protects colonic epithelial cells from acute heme-induced stress through heme catabolism; in its absence, persistent heme triggers a compensatory stress-adaptive transcriptional program and cell cycle arrest. To further assess iron-induced oxidative damage in response to heme, we performed flow cytometry for the lipid peroxidation byproduct 4-hydroxynonenal (4-HNE) in control and KO colonoids exposed to hemin. KO colonoids exhibited significantly reduced 4-HNE levels (median fluorescence intensity) and percent positive cells compared to controls (Fig. 1M), suggesting that heme-iron released by HO-1 contributes to lipid peroxidation in colonic epithelial cells. These results reinforce the role of HO-1 in regulating oxidative stress responses and susceptibility to lipid damage. However, increased cell death in the KO colonoids points to non-oxidative injury likely induced by ongoing heme stress.

### HO-1 expression is induced alongside glutathione and ferroptosis response pathways in murine and human colonic tissue and organoids

Given the activation of glutathione metabolism and ferroptosis-response pathways in HO-1-deficient colonoids in response to heme (Fig. 1), we examined whether similar stress responses are activated in colonic epithelial cells in response to induction of ferroptosis and inflammation. WT murine colonoids were exposed to erastin, a glutathione-depleting compound that induces oxidative stress and eventually ferroptosis by inhibiting the cystine/glutamate antiporter system X_C-_ encoded by *Slc7a11* and *Slc3a2* (18). There was a significant upregulation of ferroptosis-response and glutathione metabolism genes, along with increased *Hmox1* expression (Fig. 2A). Similarly, human colonoids derived from ulcerative colitis (UC) patient biopsies exhibited higher baseline and erastin-induced expression of ferroptosis and glutathione metabolism genes compared to colonoids from healthy control biopsies (Fig. 2B-C). This includes *HMOX1*, suggesting that HO-1 is consistently upregulated in epithelial cells under pro-ferroptotic conditions.

**Figure 2.**
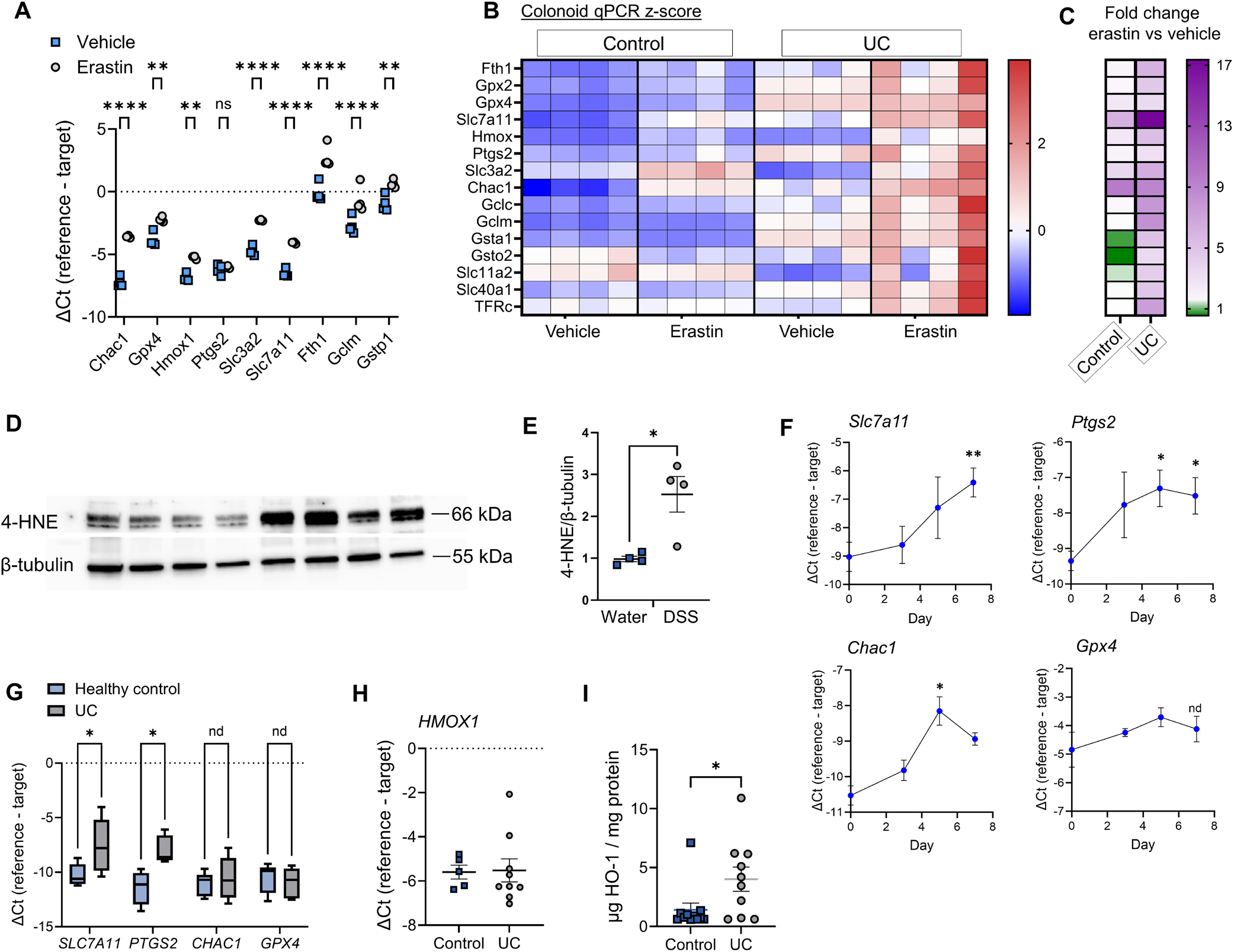
HO-1 is induced alongside ferroptosis and antioxidant responses in murine and human colonic tissue and organoids. (A) Murine colonoid mRNA expression by RT-qPCR after exposure to vehicle or erastin (20µM) for 24h. (B) Z-score of gene expression in healthy human vs UC-derived colonoid by RT-qPCR after exposure to vehicle or erastin (20µM) for 24h. (C) Log2 fold change reflects comparison between erastin and vehicle treated groups within each cell type only. (D-E) WT mice were given DSS 2.5% in their drinking water for 5 days and colonic tissue was collected and analyzed by Western blot for ferroptosis marker 4-HNE compared to mice only given drinking water alone. Levels of 4-HNE were calculated relative to β-tubulin (n =4). (F) Whole colon mRNA expression by RT-qPCR of tissue from WT mice that were given DSS 2.5% in their drinking water for 5 days followed by drinking water alone for 2 days after DSS was removed (n = 3-4 mice per time point). (G) Human colonic biopsy tissue mRNA expression by RT-qPCR (n = 4-6 control, 7-9 UC). (H) *HMOX1* mRNA from human colonic biopsy tissue. (I) Human colonic biopsy tissue HO-1 assessed by ELISA of tissue homogenates relative to total protein. Data represent mean ± SEM; \**P* < 0.05, \*\**P* < 0.01, \*\*\*\**P* < 0.0001 by unpaired Student’s *t* test, multiple *t* test for two groups / ANOVA for three or more groups with correction for multiple comparisons.

Lipid peroxidation has recently been identified as a feature of intestinal inflammation (19). We assessed the physiological relationship of HO-1 induction seen in DSS colitis (Fig. 1) to markers of ROS-mediated lipid injury in vivo. During the course of DSS injury, we observed increased levels of 4-HNE (Fig. 2D-E), along with increased expression of well-reported marker genes including *Ptgs2, Slc7a11, and Chac1* (Fig. 2F) (20). Gene expression in UC biopsy tissue showed similar trends with increased ferroptosis responsive genes such as *PTGS2* and *SLC7A11* (Fig. 2G). We examined a publicly available Gene Expression Omnibus data set (GSE38713) that reported transcriptional patterns in colon tissue from healthy individuals and from UC patients (32). A similar elevation of ferroptosis-linked gene expression was seen, particularly in actively inflamed tissue from UC patients, similar to changes seen at the peak of DSS injury on day 7 (Fig. S2). In our own tissue samples from healthy individuals and those with UC, we detected increased expression of HO-1 protein consistent with prior studies in UC patients (33-35), while *HMOX1* mRNA remained unchanged suggestive of post-transcriptional regulation (Fig. 2H-I). Together, these results demonstrate that HO-1 induction correlates with activation of glutathione metabolism pathways and intersects with the 0ferroptosis-response pathway in the context of experimental and disease-associated inflammation and oxidative stress.

### HO-1 and ferroptosis markers are upregulated in colonic tumors and correlate with lipid peroxidation

Chronic colitis is a well-established risk factor for colorectal cancer, primarily due to persistent oxidative stress and cycles of epithelial injury and repair. Emerging evidence implicates lipid peroxidation in the pathogenesis of tumorigenesis (36). Developing tumors often exhibit enhanced resistance to oxidative stress, facilitated by upregulation of antioxidative pathways that support cancer stem cell survival and proliferation (37). Paradoxically, colorectal cancer cells also exhibit a heightened demand for iron to sustain metabolic activity and growth, despite the risk of iron-induced oxidative damage (22, 38).

To investigate the role of HO-1 in this context we employed the azoxymethane-dextran sodium sulfate (AOM-DSS) model of colitis-associated cancer (Fig. 3A), which models chronic epithelial injury leading to the development of colonic tumors (39). Using WT mice, we observed significantly increased expression of ferroptosis-response genes in the colon tumors, compared to adjacent uninvolved colon, including *Hmox1, Ptgs2* and *Slc7a11* (Fig. 3B). These findings suggest that the tumor microenvironment (TME) is characterized by elevated oxidative stress and lipid peroxidation.

**Figure 3.**
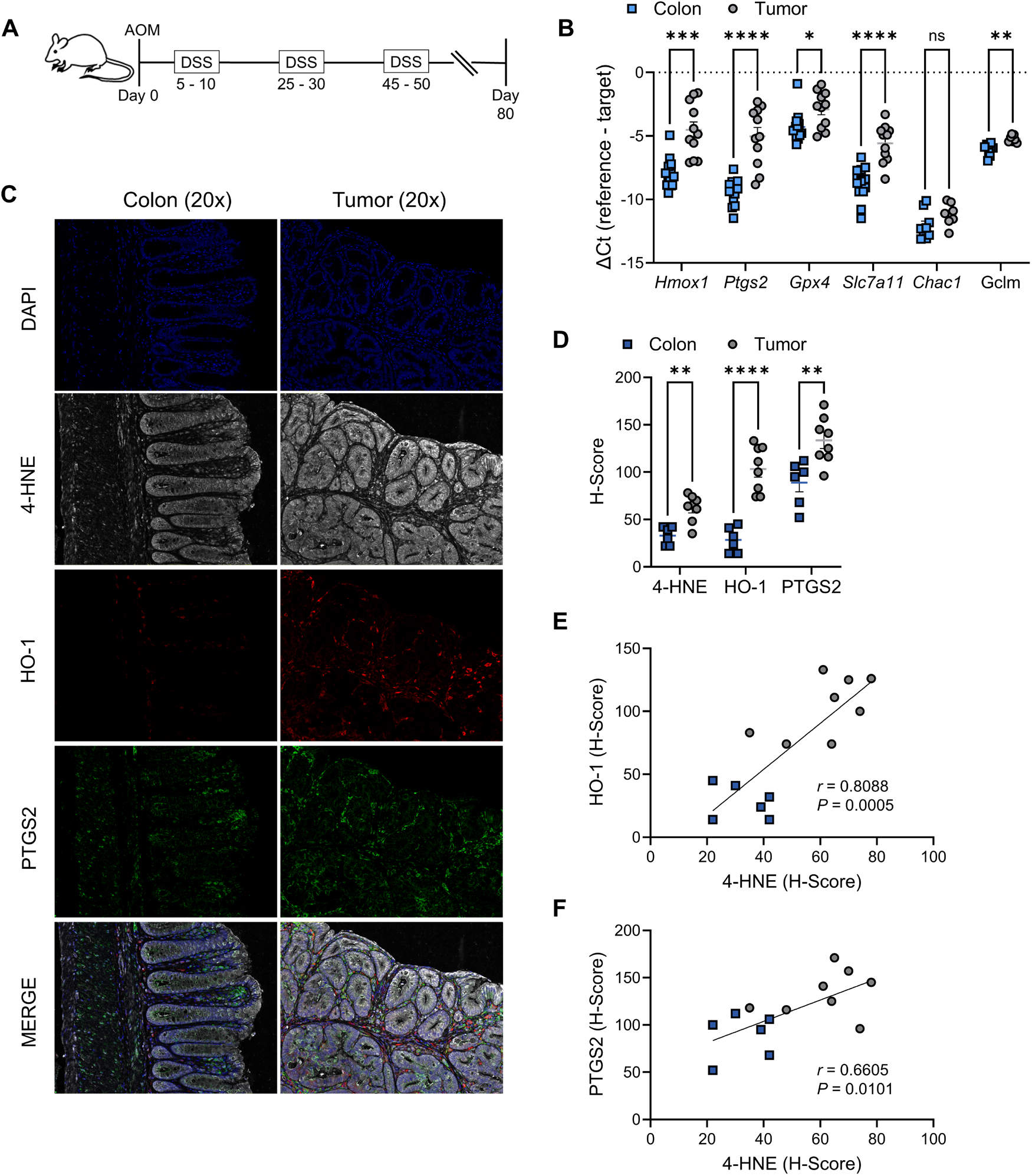
HO-1, lipid peroxidation and ferroptosis response genes are upregulated in colonic tumors. (A) Scheme of method for azoxymethane-DSS colitis-associated cancer model. (B) Gene expression by RT-qPCR of tissue from individually pooled tumors from 14 mice, and adjacent matched uninvolved colon tissue from 9-14 mice via AOM-DSS model in WT C57BL/6 mice. (C) Representative photomicrographs of immunofluorescent staining using paraffin-embedded mouse tumor and adjacent colon sections from an AOM-DSS colitis experiment. (D) Histologic scores of proteins from individual tumor sections from 8 mice and adjacent colon sections from 6 mice. (E-F) Pearson correlations of the lipid peroxidation byproduct 4-HNE against HO-1 or PTGS2. Data represent mean ± SEM; \**P* < 0.05, \*\*\**P* < 0.001, and \*\*\*\**P* < 0.0001, by multiple unpaired *t* test with correction for multiple comparisons (colon vs tumor for each gene.

To validate these transcriptional changes at the protein level, we performed quantitative multiplexed immunofluorescence using the PhenoImager HT platform. We observed significantly increased levels of PTGS2, 4-HNE and HO-1 in tumor sections compared to matched non-tumor colon tissue (Fig. 3C-D). Notably, 4-HNE levels strongly correlated with detection of HO-1 and PTGS2, reinforcing the link between HO-1 induction and oxidative lipid damage within the tumor microenvironment (TME; Fig. 3E-F). Collectively, these data support increased lipid peroxidation in the TME while the consistent and strong association with HO-1 suggests it could be a biomarker of oxidative stress activity in colitis-associated tumors and warrants further investigation of a potential mechanistic role.

### Deletion of epithelial HO-1 reduces tumor burden and colitis-associated tumorigenesis

Given the consistent upregulation of HO-1 in colonic tumors and its association with oxidative stress, we next investigated whether epithelial HO-1 plays a functional role in colitis-associated tumorigenesis. Using the AOM-DSS model, we compared tumor development in our control *Hmox1*^fl/fl^ mice and our *Hmox1*^ΔIEC^ mice during the course of chronic colitis. We observed similar weight loss, bleeding scores and fecal heme content between the two groups (Figs. 4A; S3). However, *Hmox1*^ΔIEC^ mice developed significantly fewer colonic tumors compared to controls (Fig. 4B-D). Tumors in knockout mice were predominantly located in the distal colon, consistent with known injury patterns in this model (Figs. 4B; S4A) (25). Both adenomas and adenocarcinomas were observed (Figs. 4C; S4B). Tumor burden, measured by total surface area and estimated volume, was significantly reduced in *Hmox1*^ΔIEC^ mice, and large tumors (>3 mm) were less frequent (Figs. 4E-F; S4C-D). The number of colonic tumors were similarly decreased in both male and female *Hmox1*^ΔIEC^ mice (Fig. S4E). Importantly, histologic injury scores during DSS colitis and at the endpoint were comparable between genotypes, indicating that reduced tumor burden was not due to differences in inflammation severity (Fig. 4G; S3E).

**Figure 4.**
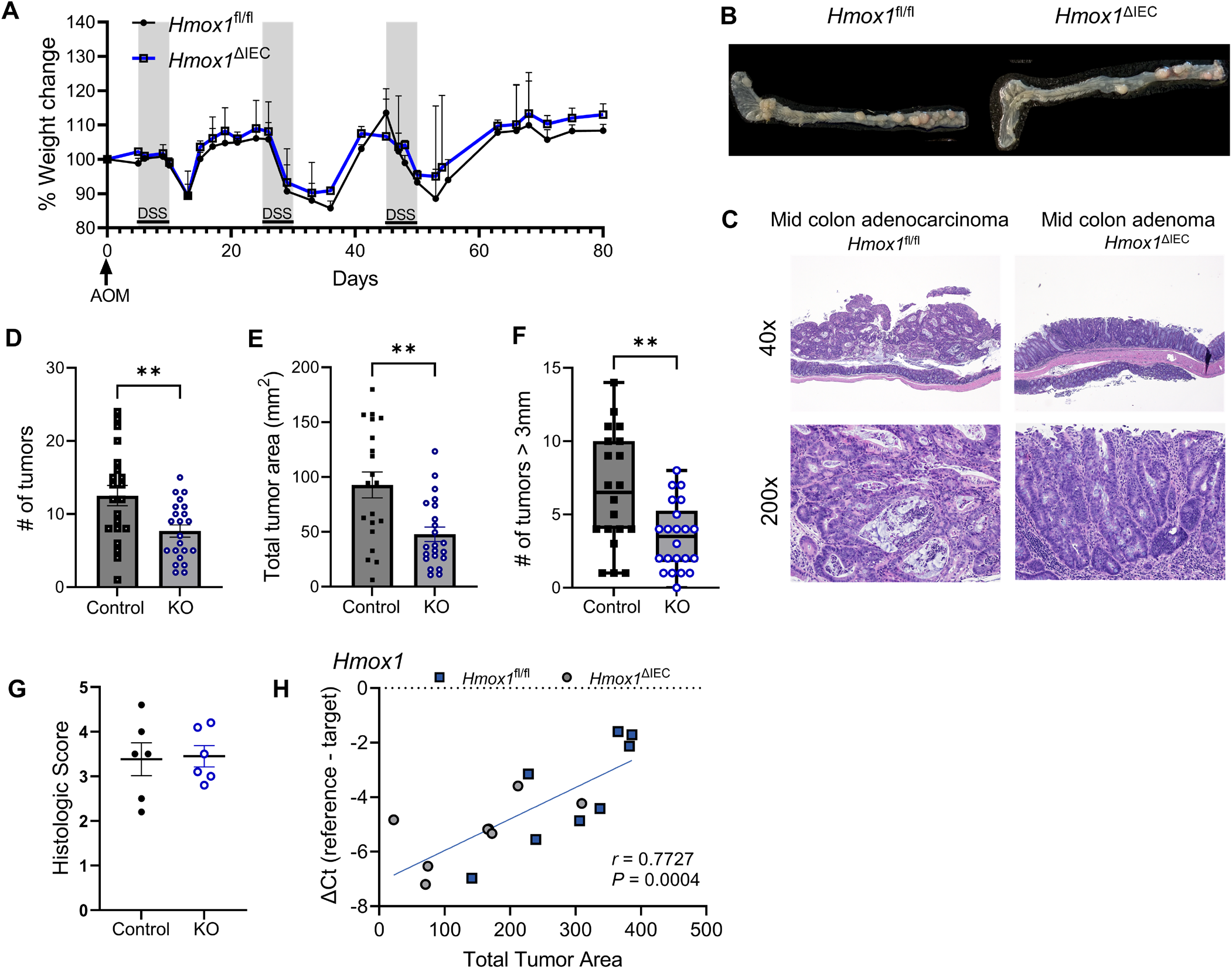
Epithelial HO-1 deletion reduces tumor burden in colitis-associated cancer. (A) Comparison of body weight changes over course of AOM-DSS colitis model in *Hmox1*^fl/fl^ and *Hmox1*^ΔIEC^ mice (n = 8-11 mice per group, pooled from 2 independent experiments). (B) Representative colons with demonstration of tumor formation at day 80 from both mouse strains. (C) Representative H&E staining of sections of colonic tumors from the AOM-DSS model. (D) Visible tumor numbers in colon + cecum at day 80 upon gross examination. Combined data from three independent experiments. (E) Tumor burden and average size as measured by 2-dimensional area of tumor involvement. (F) Number of tumors > 3mm in diameter visualized in colon + cecum at day 80 upon gross examination. (G) Colonic histopathologic injury score at day 80. (H) Pearson correlation of pooled whole tumor *Hmox1* mRNA expression (qPCR) with tumor burden. Data represent mean ± SEM; \**P* < .05, \*\**P* < .01, by unpaired, Student’s *t* test.

To further explore the relationship between HO-1 and tumorigenesis, we analyzed *Hmox1* mRNA expression in colonic tissue and assessed its correlation with tumor burden across both control and knockout mice. A strong positive correlation was observed between *Hmox1* expression and total tumor area and volume, supporting the hypothesis that elevated HO-1 may be associated with enhanced tumor growth (Fig. 4H). Notably, *Hmox1* expression levels in knockout mice were not uniformly the lowest, likely reflecting contributions from non-epithelial cell types such as immune or stromal cells, suggesting further complexity in cell-type-specific roles for HO-1 in tumorigenesis.

Collectively, these results provide functional evidence that epithelial HO-1 influences tumor development in colitis-associated cancer. Given prior findings linking HO-1 to regulation of oxidative stress responses, we next examined the consequences of HO-1 deletion on oxidative damage within the TME.

### Epithelial HO-1 regulates iron availability and tumor epithelial transcriptional adaptation to oxidative stress

To investigate the impact of epithelial HO-1 deletion on oxidative stress and its influence on tumor epithelial proliferation in our CAC model, we stained for 8-hydroxy-2’-deoxyguanosine (8-OHdG), which measures oxidative DNA damage and Ki-67, a marker of proliferation, alongside DAPI and EpCAM. Tumors from knockout mice exhibited significantly reduced 8-OHdG staining compared to controls (Fig. 5A-B). Interestingly, this reduction was also observed in uninvolved colonic tissue (Fig. 5C), suggesting that epithelial HO-1 contributes to oxidative DNA damage in chronically inflamed and neoplastic tissue. Ki-67 staining revealed a modest, non-significant reduction in the percentage of proliferating cells in knockout tumors (Fig. 5D). However, Ki-67 expression levels, as measured by H-score, were significantly lower in *Hmox1*^ΔIEC^ tumors (Fig. 5E), indicating reduced proliferative activity. These findings support a role for epithelial HO-1 in promoting tumor cell proliferation in CAC, possibly by regulating oxidative stress and damage to DNA.

**Figure 5.**
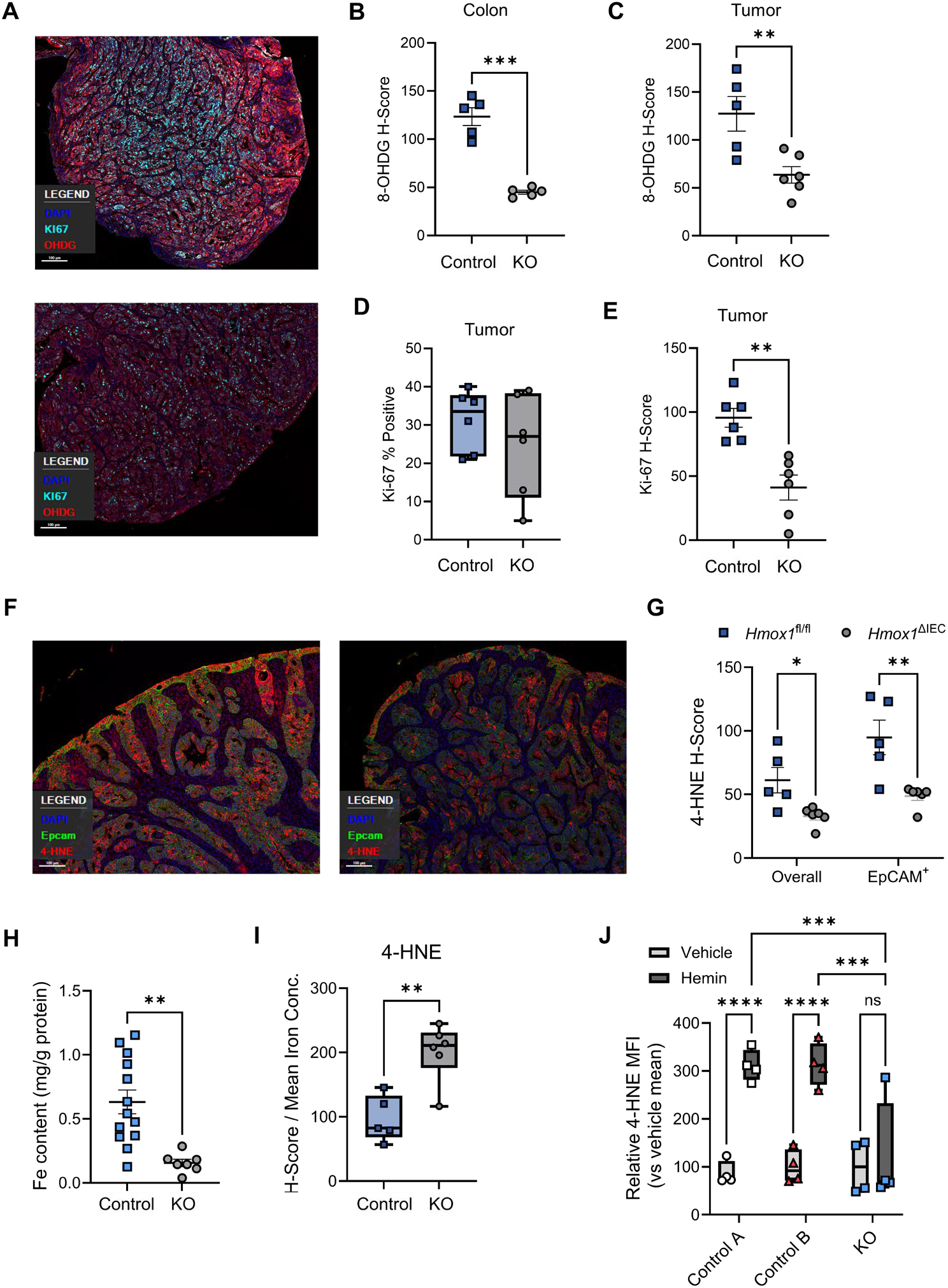
Epithelial HO-1 regulates oxidative damage, iron levels, and stress adaptation in the TME. (A) Representative photomicrographs of immunofluorescent staining for 8-hydroxy-2’-deoxyguanosine (8-OHdG) and Ki-67, using paraffin-embedded mouse colon individual tumor sections from 4 control and 4 *Hmox1*^ΔIEC^ mice undergoing AOM-DSS. (B) Quantification of 8-OHdG staining intensity in tumor tissue. (C) 8-OHdG staining intensity in uninvolved colonic tissue from control and knockout mice. (D) Percentage of Ki-67+ cells based on immunofluorescence staining. (E) Ki-67 H-score analysis from tumor sections. (F) Representative photomicrographs of immunofluorescent staining for 4-HNE and EpCAM using colon tumor sections. (G) Quantification of 4-HNE from tumor sections taken from control and knockout mouse tumors. (H) Whole tumor iron concentration measured by colorimetric assay. (I) 4-HNE staining intensity normalized to mean tumor iron levels per genotype. (J) Tumoroids derived from pooled tumors from 2x *Hmox1*^fl/fl^ (control A and B) and 1x *Hmox1*^ΔIEC^ (KO) mice were exposed to hemin (100 µM) for 24h. Lipid peroxidation measured by flow cytometry for 4-HNE with median fluorescence intensity (MFI) values normalized to the mean of each vehicle-treated group. Data in figures represent mean ± SEM; \**P* < .05, \*\**P* < .01, \*\*\**P* < .001 by unpaired, Student’s *t* test, multiple unpaired *t* test for two groups / one-way ANOVA for three or more groups with correction for multiple comparisons.

To explore the relationship between HO-1 deletion and lipid peroxidation, we quantified 4-HNE staining in tumor sections. Representative images and H-score analysis of whole tumors and EpCAM+ epithelial cells showed significantly reduced 4-HNE staining in tumors from *Hmox1*^ΔIEC^ mice compared to controls (Fig. 5F-G). A previous study demonstrated that overexpression of HO-1 in a CRC cell line increases the labile iron pool and promotes lipid peroxidation (40). We therefore measured tumor iron content to determine whether loss of epithelial HO-1 influences lipid peroxidation via regulation of iron levels. We measured significantly decreased iron in tumors from *Hmox1*^ΔIEC^ mice (Fig. 5H), consistent with expectations that impaired heme catabolism can reduce labile iron levels and influence iron-dependent oxidative damage. Paradoxically, when tumor 4-HNE staining was normalized to average (mean) iron content by strain, knockout tumors showed significantly higher 4-HNE per unit iron (Fig. 5I), suggesting increased iron-independent oxidative lipid damage. This could reflect oxidative damage driven by uncatabolized heme, rather than free iron, consistent with known oxidative and non-oxidative mechanisms of heme toxicity (24, 41).

As colonic tumors grow, increased and often dysregulated vascularity can lead to microhemorrhages and cell death, resulting in elevated levels of heme, hemoglobin, and hemeproteins (42). To further validate the influence of epithelial HO-1 in regulating tumor epithelial responses to heme within the TME, as a proof-of-concept, we developed tumor epithelial-derived organoids (tumoroids) from two control (*Hmox1*^fl/fl^) and one KO (*Hmox1*^ΔIEC^) mouse. After exposure to hemin, control tumoroids demonstrated significantly elevated 4-HNE levels compared to vehicle-treated controls, consistent with HO-1-mediated iron release causing lipid peroxidation (Fig. 5J). In contrast, KO tumoroids showed no increase in 4-HNE after exposure to hemin (Fig. 5J), supporting a requirement for HO-1 in hemin-induced lipid peroxidation in tumor epithelial cells. These results reinforce the role of HO-1 in shaping epithelial oxidative stress responses in the TME.

### Single-cell transcriptomic profiling reveal stress-adaptive epithelial responses

Given the potential for non-catabolized heme to induce cellular stress independent of iron, we further explored how loss of HO-1 influences epithelial cell stress responses within the TME by performing single-cell RNA sequencing (scRNA-seq). Pooled tumors from *Hmox1*^fl/fl^ and *Hmox1*^ΔIEC^ mice > 2mm in diameter were enzymatically dissociated, and live cells were sorted for sequencing using the 10X Genomics platform (Fig. 6A). After quality control, library normalization and dimensionality reduction, ∼24,000 cells were co-embedded in UMAP space (Fig. 6B). Unsupervised clustering initially identified 18 distinct populations, which were annotated using a combination of canonical marker genes (e.g., T cells: *CD3e*; B cells: *CD79a*; tumor epithelial cells: *Epcam*; neutrophils: *Cxcr2*; monocyte/macrophages: *Fcgr1*; mast cells: *Fcer1a*), and guidance from prior scRNA-seq datasets from AOM-DSS and murine intestine (Figs. 6C; S5) (43-45). Both genotypes exhibited similar cellular composition, including tumor epithelial cells and diverse leukocyte populations (Fig. 6B).

**Figure 6.**
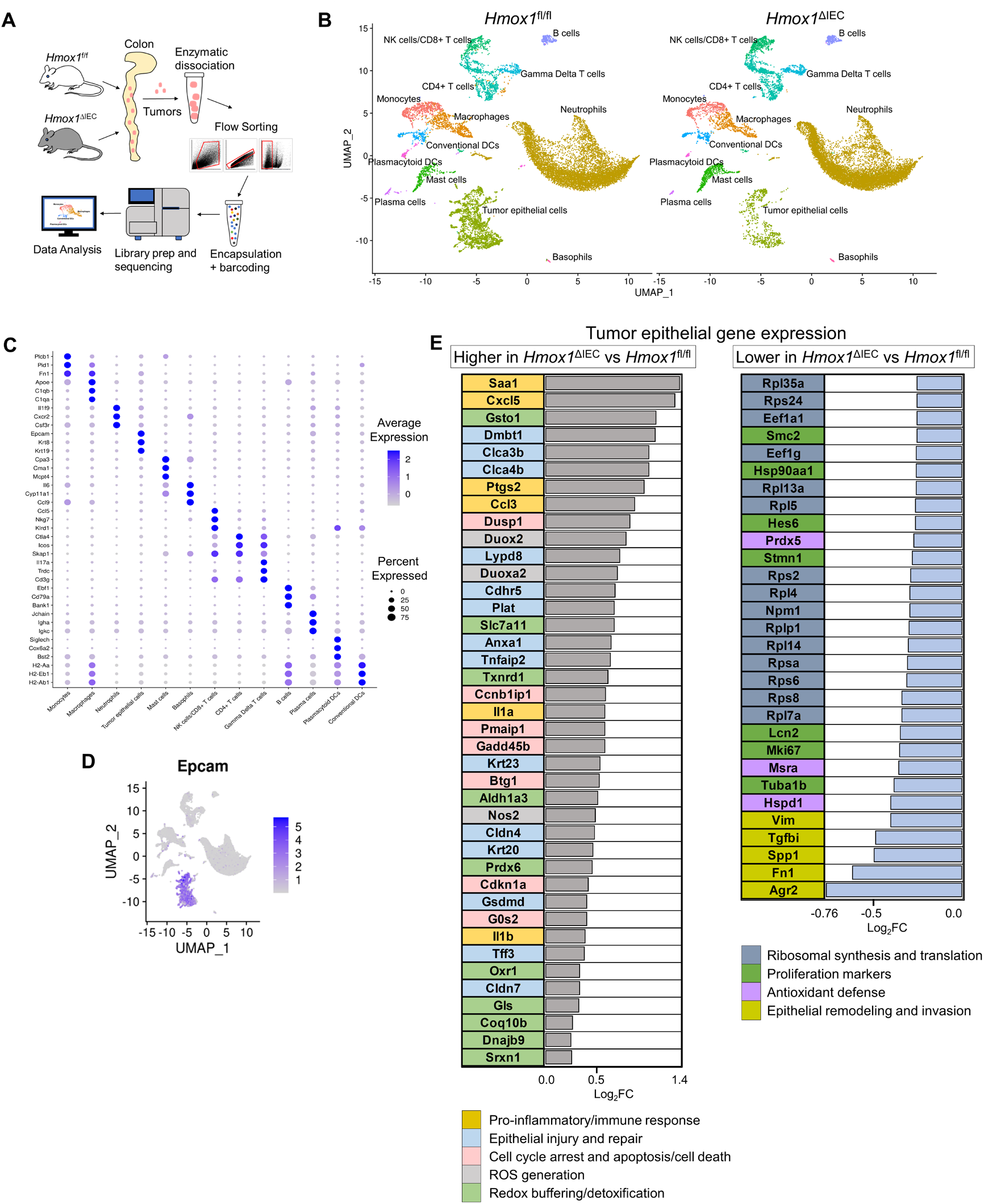
Loss of epithelial HO-1 induces a stress-adaptive transcriptional shift in tumor epithelial cells. (A) Overview for single-cell RNA sequencing workflow. Pooled tumors from control and knockout mice were enzymatically dissociated and live cells sorted for scRNA-seq using the 10x Genomics platform. (B) UMAP plot of ∼24,000 quality-controlled cells from both genotypes, showing unsupervised clustering and manual annotation of major cell types. (C) Dot plot of top three marker genes per cluster used to validate annotations, ranked according to log_2_ fold change multiplied by difference in percent expression compared to all other cells. (D) Feature plot of *Epcam* expression across the UMAP, highlighting the epithelial subcluster used for differential expression analysis. (E) Bar graphs of significantly upregulated and downregulated genes in *Epcam*+ epithelial cells from knockout tumors, identified using the MAST hurdle model with Bonferroni-adjusted *P* < 0.05 and log_2_FC > 0.25. Genes are color-coded by functional annotation.

Analysis of the *Epcam*+ tumor epithelial cell cluster (Fig. 6D) revealed distinct transcriptional programs between genotypes (Fig. 6E). In knockout epithelial cells, a reactive and stress-adaptive phenotype was observed, marked by oxidative stress responses, heightened inflammation, cell cycle regulation, and stress-induced programmed cell death. This profile is driven by a robust oxidative stress response, with increased expression of genes involved in ROS generation (*Duox2, Duoxa2, Nos2*) and redox buffering/detoxification (*Txnrd1, Gsto1, Aldh1a3, Slc7a11, Prdx6, Oxr1, Dnajb9*), consistent with efforts to mitigate oxidative damage and maintain cellular homeostasis. This transcriptional shift is consistent with an adaptive response to toxicity from uncatabolized heme and aligns with the stress-adaptive phenotype observed in vitro (24). In line with this, genes regulating cell cycle arrest and apoptosis (*Cdkn1a, Ccnb1ip1, Gadd45b, Btg1, Pmaip1, Gsdmd, Dusp1*) were elevated, reflecting a tightly controlled response to cellular stress. Elevated expression of genes involved in pro-inflammatory and immune responses (*Saa1, Cxcl5, Ccl3, Il1a, Il1b, Ptgs2*) was also observed. Genes associated with epithelial injury and repair (*Dmbt1, Lypd8, Clca3b, Clca4b, Plat, Anxa1, Tnfaip2, Krt20, Krt23, Cldn4, Cdhr5*) were also upregulated, suggesting active tissue repair/remodeling.

In contrast, control epithelial cells exhibited a transcriptional profile consistent with a more aggressive, proliferative, and invasive tumor microenvironment. These cells upregulated genes linked to iron metabolism and metastatic potential (*Lcn2*), epithelial remodeling and invasion (*Tgfbi, Spp1, Agr2, Fn1, Vim*), and general antioxidant defense (*Prdx5, Msra, Hspd1*). Elevated expression of proliferation markers such as *Mki67, Hsp90aa1, Tuba1b* and *Hist1h4c* align with increased Ki-67 staining, supporting a more proliferative tumor phenotype.

## Discussion

Colitis-associated colorectal cancer (CAC) arises in the context of chronic mucosal inflammation, epithelial injury, and hemorrhage, which together create a pro-oxidant microenvironment rich in luminal heme. While epithelial HO-1 is traditionally viewed as cytoprotective, our findings reveal a context-dependent role in tumorigenesis. Specifically, epithelial HO-1 protects against *acute* heme toxicity (10, 14) but paradoxically promotes tumor growth in *chronic* inflammation by increasing intracellular iron availability and amplifying oxidative damage.

Our data demonstrate that HO-1 is consistently upregulated in colonic epithelium during active colitis and in tumors, correlating with measures of oxidative stress (33-36). Beyond mechanisms driven by iron released through HO-1-mediated catabolism, intact heme itself can exert cytotoxic effects through multiple pathways. These include intercalating into lipid membranes, proteasome inhibition, disrupting membrane integrity, generating ROS without iron release, and damaging intracellular organelles via both oxidative and non-oxidative mechanisms (24, 41, 46). These alternative modes of heme-induced injury may be particularly relevant in the absence of HO-1, which normally facilitates heme detoxification through catabolism and iron sequestration. The cytoprotective role of HO-1 under acute heme stress was evident in our colonoid experiments where heme exposure to HO-1-deficient colonoids led to increased cell death, cell cycle arrest, and a robust stress-adaptive transcriptional response designed to compensate for lost antioxidative and detoxification function (10, 30, 31). In chronically inflamed colonic epithelium of *Hmox1*^ΔIEC^ mice this compensatory response, along with regulation of iron availability, may influence tumor initiation by reducing oxidative DNA damage. In line with this, in *Hmox1*^ΔIEC^ mice, we observed a significant reduction in tumor number and size, despite comparable levels of colitis-induced injury. This suggests that epithelial HO-1 contributes directly to tumorigenesis, independent of inflammation severity, in contrast to many studies on CAC (39). Mechanistically, HO-1 deletion led to reduced oxidative DNA damage (8-OHdG), diminished 4-HNE accumulation as a product of lipid oxidation, and lower iron levels in tumors (36, 40). These findings also support a model in which epithelial HO-1-mediated heme catabolism augments tumor growth by increasing bioavailable iron, fueling oxidative stress and damage but more importantly, driving proliferation (15, 22, 23). Iron promotes tumor growth by supporting nucleotide metabolism and proliferation (22, 23), positioning HO-1 as a key upstream regulator of iron-dependent metabolic reprogramming in the TME. In its absence, reduced iron availability and the compensatory stress response may jointly suppress tumor progression. Interestingly, when 4-HNE was normalized to iron content, knockout tumors exhibited higher 4-HNE per unit iron. This suggests a less potent source of oxidative damage, independent of free iron, likely due to uncatabolized heme (24).

Single-cell RNA sequencing revealed that tumor epithelial cells from *Hmox1*^ΔIEC^ mice adopt a stress-adaptive transcriptional program characterized by upregulation of redox buffering, cell cycle arrest, and programmed cell death pathways. This stress-adaptive phenotype, likely driven by persistent heme toxicity, may suppress tumor progression by limiting proliferation and promoting repair. In contrast, control tumor epithelial cells exhibited a more proliferative and invasive phenotype, with elevated expression of iron metabolism genes and markers of epithelial-mesenchymal transition. These transcriptional differences mirror the histologic and molecular phenotypes, reinforcing HO-1’s role in promoting a proliferative tumor cell state.

Our findings contribute to the growing literature on iron-dependent oxidative stress and lipid peroxidation in gastrointestinal cancers. Ferroptosis, driven by iron-dependent lipid peroxidation, represents one potential outcome of dysregulated iron metabolism, and HO-1 has been reported to both promote and suppress ferroptosis, depending on cellular context and iron availability (47-52). However, our data primarily support a contextual role for HO-1 in regulating oxidative damage and proliferation through regulation of iron availability. In colonic epithelial cells exposed to heme, HO-1 appears protective, limiting cell death and oxidative damage (10, 31). However, in the tumor microenvironment, HO-1 may facilitate iron accumulation and oxidative stress, promoting proliferation and tumor progression (22, 40). This duality has important implications. Therapeutic strategies targeting HO-1 or iron metabolism must consider the temporal and spatial context of HO-1 activity. Inhibiting HO-1 may sensitize tumor cells to ferroptosis or constrain proliferation by limiting iron availability, but could also impair epithelial resilience to acute heme stress (53). Conversely, enhancing HO-1 activity may protect against inflammation-induced damage but risk promoting tumorigenesis in chronic disease.

In summary, our study identifies epithelial HO-1 as a key regulator of iron metabolism, oxidative stress, and tumor epithelial cell adaptation in colitis-associated cancer. The apparent context-dependent function of HO-1, cytoprotection in acute injury but tumor-promotion in chronic inflammation, highlight its complex role in colonic epithelial homeostasis and neoplasia. Future studies could further explore the therapeutic implications of manipulating epithelial HO-1 and lipid peroxidation in other models of IBD and colorectal cancer.

## Methods

### DSS colitis vs Azoxymethane-DSS colitis

C57BL/6J WT mice were used where indicated and C57BL/6 *Hmox1*^fl/fl^ mice were kindly provided by Dr. H. B. Suliman (54). *Hmox1*^ΔIEC^ mice were made by crossing *Hmox1*^fl/fl^ mice with B6.Cg-Tg(Vil1-cre)1000Gum/J mice expressing Cre recombinase in villus and crypt epithelial cells of the small and large intestines (JAX #021504) (55). All mice were maintained and managed in accordance with Institutional Animal Care and Use Guidelines. Littermate controls were used for all colitis experiments. We used age-matched male and female mice starting between age 7 and 14 weeks. For colitis experiments, mice were given 2.5% colitis grade dextran sodium sulfate (DSS, MP Biomedical) in drinking water, changed every 2 days, for 5 days and were then sacrificed at day 7 where indicated. For the colitis-associated cancer model, a single dose of AOM at 10 mg/kg of body weight was administered intraperitoneally on day zero and five days later, mice underwent three rounds of five days of 2.5% DSS in drinking water, followed by 14-days of tap water, with the final period of tap water extended to 30 days (Fig. 3A). Mice were sacrificed on day 80. Disease activity components were measured blindly (weight change, stool consistency, and hematochezia scores; scale 0-3) and an index was computed by summing the scores (max score of 9). At experiment end, colons were excised, cleaned, the lumen was then exposed, and photos were taken with ruler or calipers for measurements of the tumors. Final tumor measurements (e.g. area) were calculated using ImageJ (56). Colons were fixed, embedded, and processed for histology. H&E stained tissue was scored by a histo-pathologist blinded to the treatments and groupings of animals using described methods (25). *Sex as a biological variable*: Our study examined male and female mice, and similar findings are reported for both sexes.

### IEC/leukocyte isolation and flow cytometry

Isolation of tumor cells and intestinal epithelial cells was performed as previously described (57). Modifications include retention of the dissociated epithelial fraction where necessary for fluorescence activated cell sorting (FACS) and measurement of *Hmox1* expression. We performed flow cytometry as previously described (57). Dead cells were stained and excluded using BioLegend 7AAD viability staining solution or Zombie Aqua™ dye prior to flow cytometry or FACS. Anti-4-HNE (R&D Systems) was used with an AF647 secondary antibody (Invitrogen) to detect lipid peroxidation via flow cytometry, which was validated using H_2_O_2_ as a positive control. Other anti-mouse antibodies were purchased from BioLegend or Thermo Fisher Scientific: CD45 (30-F11); EpCAM (G8.8). Flow cytometry was performed on a FACSCanto™ II (Becton Dickinson), or a BD FACSAria™ Fusion and resulting data was analyzed with De Novo’s FCS Express 7 software.

### Cell culture

Epithelial cells were harvested from 8 to 12-week-old *Hmox1*^fl/fl^ and *Hmox1*^ΔIEC^ mouse colons or tumors, and processed as previously described to make colonoids (58). Tumor-derived organoids were made from pooled tumors from two control mice separately and one knockout mouse. Data is representative of 2-3 independent experiments with identical results. Human colonic epithelial organoids were obtained by biopsy via colonoscopy. Media was supplemented L-WRN and with ROCK inhibitor Y-27632. Organoids were cultured for 7-10 days, passaged and used typically 48h after plating. Organoids were also frozen and used after successful thawing. Hemin was purchased from Sigma-Aldrich. Erastin was purchased from MedChemExpress.

### Reverse transcription-quantitative PCR (qPCR) and mRNA sequencing

RNA was isolated from snap frozen tissue samples or cells using Trizol (Invitrogen) or Qiagen RNeasy Mini kit respectively, with the addition of the gDNA-eliminator columns as per manufacturer’s protocol. First-strand complementary DNA synthesis was performed with using iScript reverse transcription super-mix from Bio-Rad. Real-time qPCR was performed in technical duplicates using SYBR green on Applied Biosystems QuantStudio 5 Real-Time PCR System. Due to variability in transcript detection across samples, the number of biological replicates included in an analysis may differ by gene. β-Actin (*Actb*) was used as the reference gene and results are reported as ΔCt, which is the Ct of the target gene subtracted from the Ct of the reference gene (higher ΔCt values = higher expression relative to the reference gene). Primer sets were purchased from Sigma-Aldrich’s KiCqStart™ line. Preparation of mRNA library, sequencing and bioinformatic analysis was conducted by Novogene Corporation Inc.

### Heme quantification assay

Total heme was measured using an established protocol with minor adaptations (59).

### Intracellular Iron Quantification

Organoids were suspended and lysed in RIPA buffer. Pierce™ BCA Protein Assay Kit was used to obtain total protein concentration of organoids. Iron Assay Kit (Colorimetric) from Novus Biologicals was used to measure total iron content. Organoid lysate was substituted for tissue homogenate in the manufacturer’s protocol. Both BCA and Iron Assay were measured using BioTek Synergy H1 microplate reader. Results were analyzed using Microsoft Excel.

### Quantitative Multiplexed Immunofluorescence

Through our collaboration with the Human Immune Monitoring Shared Resource (HIMSR) at the University of Colorado School of Medicine we performed multispectral imaging using the PhenoImager HT instrument (formerly Vectra Polaris, Akoya Biosciences). Deparaffinized FFPE tissue sections were heat treated in antigen retrieval buffer, blocked, and incubated with various primary antibodies HO-1 (Abcam; 1:1000; pH9), PTGS2 (Cell Marque; 1:50; pH6), 4-HNE (Abcam; 1:100; pH6), 8-OHdG (BioSS; 1:300; pH9) and EpCAM (1:200; pH6), followed by horseradish peroxidase-conjugated secondary antibody polymer, and HRP-reactive OPAL fluorescent reagents. Slides were stripped between each stain with heat treatment in antigen retrieval buffer. Whole slides were imaged with PhenoImager HT v2.0.0, 20x objective, 0.5-micron resolution. Images were then analyzed using inForm software v3.0 (Akoya Biosciences).

### Western Blot

Total protein content was obtained using Pierce™ BCA Protein Assay Kit. 10% Mini-PROTEAN® TGX™ Precast Protein Gel from Bio-Rad was used to separate proteins in 10% SDS-PAGE. Using Trans-Blot Turbo RTA Midi 0.2 µm PVDF Transfer Kit from Bio-Rad, proteins were transferred onto membranes in the kit. Membranes were blocked with 5% nonfat milk blocking buffer and then incubated overnight at 4° C with primary antibody 4-HNE (1:1000) from Abcam. The control used was β-tubulin (Sigma-Aldrich). Then blots were stained with secondary antibody. Protein bands were enhanced using Clarity™ Western ECL Substrate from Bio-Rad and analyzed using Bio-Rad ChemiDoc MP Imaging System.

### Human colonic biopsy tissue

Patient colon biopsy samples were collected during colonoscopy and made available as part of a biobank repository at the University of Colorado Anschutz Medical Campus Crohn’s and Colitis Center with Colorado Multiple Institutional Review Board approval. ELISA for HO-1 was performed using a human HO-1 DuoSet IC kit from R&D Systems, and total protein was assessed using a Pierce™ BCA Protein Assay Kit.

### NCBI Gene Expression Omnibus

In data set GSE38713, colonic gene expression in patients with ulcerative colitis and non-inflammatory controls was assayed using Affymetrix GeneChip high-density oligonucleotide HGU133 Plus 2.0 microarrays (32). Initial analysis was performed using GEO2R with log_2_ transformation and *P*-values were adjusted using Benjamini and Hochberg FDR method (60). Only probe sets that represent a single gene (labeled as *_at) were used.

### Single cell RNA sequencing

Pooled tumors from each of two *Hmox1*^fl/fl^ and two *Hmox1*^ΔIEC^ mice were used.

#### Processing

Processing and statistical analyses were performed in R version 4.2. (61) and Seurat version 4.3.0 (62). Seurat defaults were used unless otherwise noted. The data were restricted to features with at least 3 detected cells and low-quality cells with more than 15% mitochondrial genes or less than 300 genes were filtered out. The DoubletFinder package version 2.0.3 (63) was then used to remove multiplets. This yielded 23,698 high quality cells across all of the samples.

Library size normalization was performed with NormalizeData and the top 2000 highly variable genes (HVGs) were identified with the FindVariableFeatures variance stabilizing transformation (VST) method. Features that were repeatedly variable across datasets were then used as anchors to create an integrated dataset across samples with the FindIntegrationAnchors and IntegrateData functions. Dimensionality reduction was performed for visualization; the integrated data were scaled and centered with ScaleData and principal component analysis was performed with RunPCA to reduce the data to 30 principal components (PCs). RunUMAP was then run on the PCs.

#### Unsupervised clustering and annotation

The FindClusters algorithm was run on the PCs with a resolution of 0.45 to identify clusters in the overall cell population. Sub-clustering of epithelial (resolution=0.038) cell populations was analogously performed. The DimPlot function was then used to create UMAPs stratified by treatment group.

Positive conserved cell type markers were identified using the FindAllMarkers function with minimum detection and log_2_-fold change (FC) thresholds set to 25% and Bonferroni adjusted *P* ≤ 0.05. The markers were used to perform manual annotation and to merge and split clusters when appropriate. The marker genes were sorted by the log_2_-FC multiplied by the difference in percent of cells expressed between the cluster and all other cells. The top marker genes were then visualized with Dotplots (ggplot2 package version 3.4.0 (64)), feature plots (FeaturePlot), and heatmaps (ComplexHeatmap package version 2.12.1 (65)).

#### Gene Ontology enrichment analysis of epithelial cell populations

Epithelial sub-population differentially expressed genes (DEGs) between *Hmox1*^fl/fl^ and *Hmox1*^ΔIEC^ mice were identified using the FindAllMarkers MAST Hurdle Approach (66) and previously described thresholds. Genes were considered DE if Bonferroni-adjusted significance was less than 0.05. DEGs were then analyzed to determine whether they were enriched relative to all genes in the dataset. Genes were first annotated to gene ontology (GO) terms with the biomaRt R package (67) and genes without annotations were eliminated.

### Statistics

Unless otherwise stated, statistical significance of comparisons was performed using Prism version 9.5 and 10 (GraphPad Software). We used unpaired two-tailed Student’s *t* test for two groups, multiple unpaired *t* tests for two groups with correction for multiple comparisons using the Holm-Sidak method, or one-way ANOVA for multiple groups followed by Bonferroni or Dunnett’s post hoc test. Error bars reflect standard error of the mean unless otherwise stated. Results were considered statistically significant if *P* < 0.05.

### Study approval

All animal studies were performed in accordance with protocols approved by the University of Colorado Anschutz Medical Campus and Rocky Mountain Regional VAMC Institutional Animal Care and Use Committees.

## Data availability

Underlying data are available from the corresponding author upon request and in the Supporting Data Values XLS file.

## Author contributions

The study was designed and supervised by JCO with the support of SPC. RCC and JCC performed most of the experiments/data acquisition with contributions from GB, REMS, FM, LZ, JCO, RMN, CHTH, CAS, MEG and IMC. JCO and AWS (bioinformatics) performed data analysis and visualization with contributions from DJO, RCC, GB and SMA. Resources and funding were provided by JCO and SPC with contributions from CHTH, AT, CAS and MEG. JCO, RCC, JCC, RMN, CHTH, GB, FM, LZ and REMS contributed to methodology. Important intellectual contributions provided by SPC, SMA and IMC. Original manuscript written by JCO with contributions from RCC, JCC and AWS. Review and editing by SPC, SMA, JCO, CHTH, AWS, DJO, RCC, AT and IMC with input from all co-authors. RCC and JCC are co-first authors with RCC assigned first as the original first-author.

