## Supplemental Figures for "Epithelial HO-1 regulates iron availability and promotes colonic tumorigenesis in a context-dependent manner"

#### Supplemental Figure 1

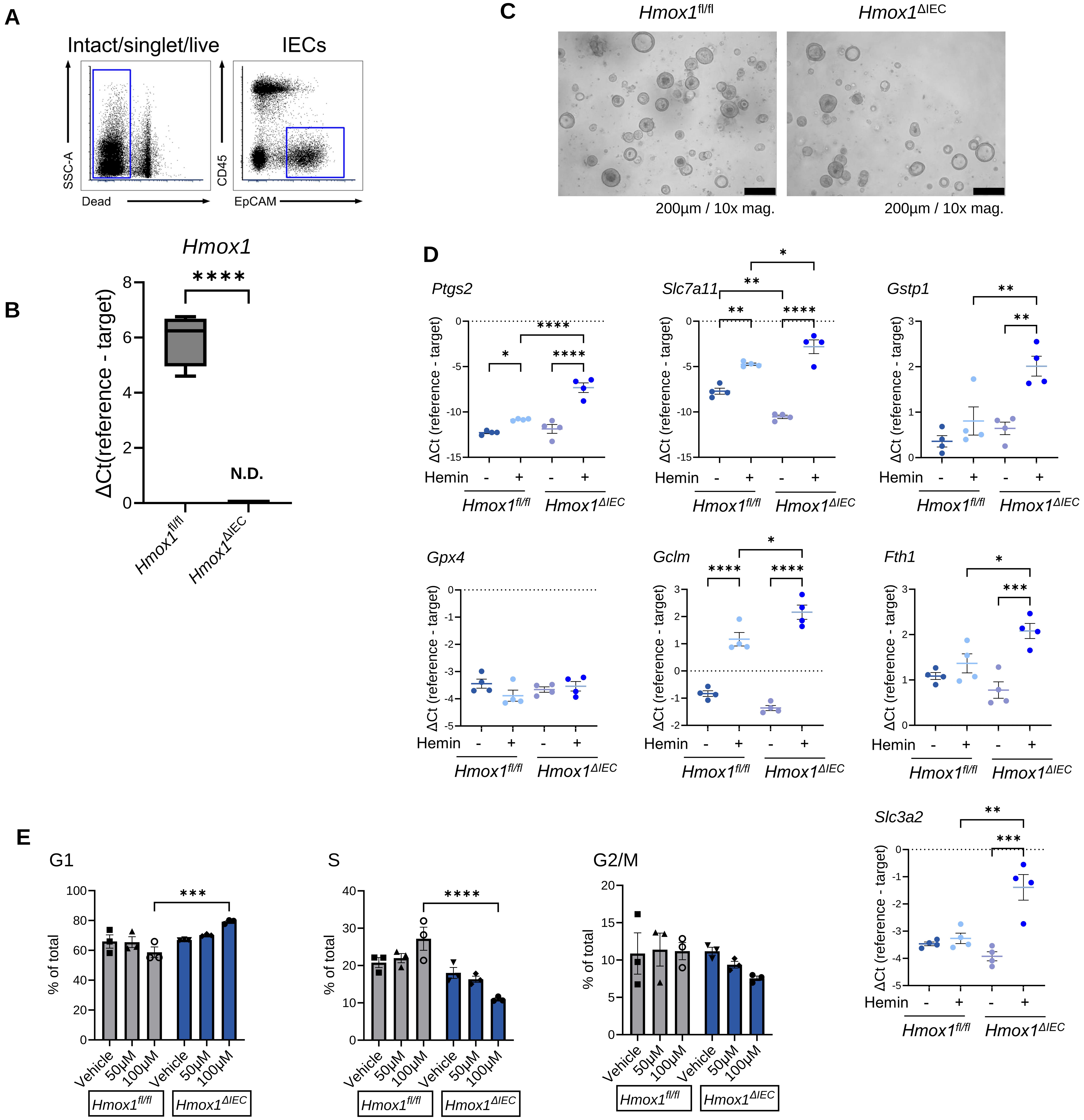


Supplemental Figure 1. *Hmox1* regulates the stress and cell cycle response to heme (A) $CD45^{-}Epcam^{+}$ colonic epithelial cells from $Hmox1^{fl/fl}$ and $Hmox1^{\Delta IEC}$ mice were sorted by FACS. Representative gating strategy shown. (B) *Hmox1* expression in colonic epithelial cells assessed using RT-qPCR (N.D. = not detected; n = 4 per group). (C) Representative photomicrographs of colonic epithelial organoids derived from healthy $Hmox1^{fl/fl}$ and $Hmox1^{\Delta IEC}$ mouse colons. (D) Murine colonic epithelial organoid mRNA expression by RT-qPCR after exposure to hemin (100µM) for 24h (n = 4 per group). (E) Cell cycle states were assessed using 7AAD and flow cytometry in murine colonic epithelial organoids exposed to hemin or vehicle for 24h in three independent experiments. Data represent mean ± SEM. **P* < 0.05, ***P* < 0.01, ****P* < 0.001, and *****P* < 0.0001, by unpaired, Student's *t* test or one-way ANOVA with correction for multiple comparisons (Bonferroni).

#### Supplemental Figure 2

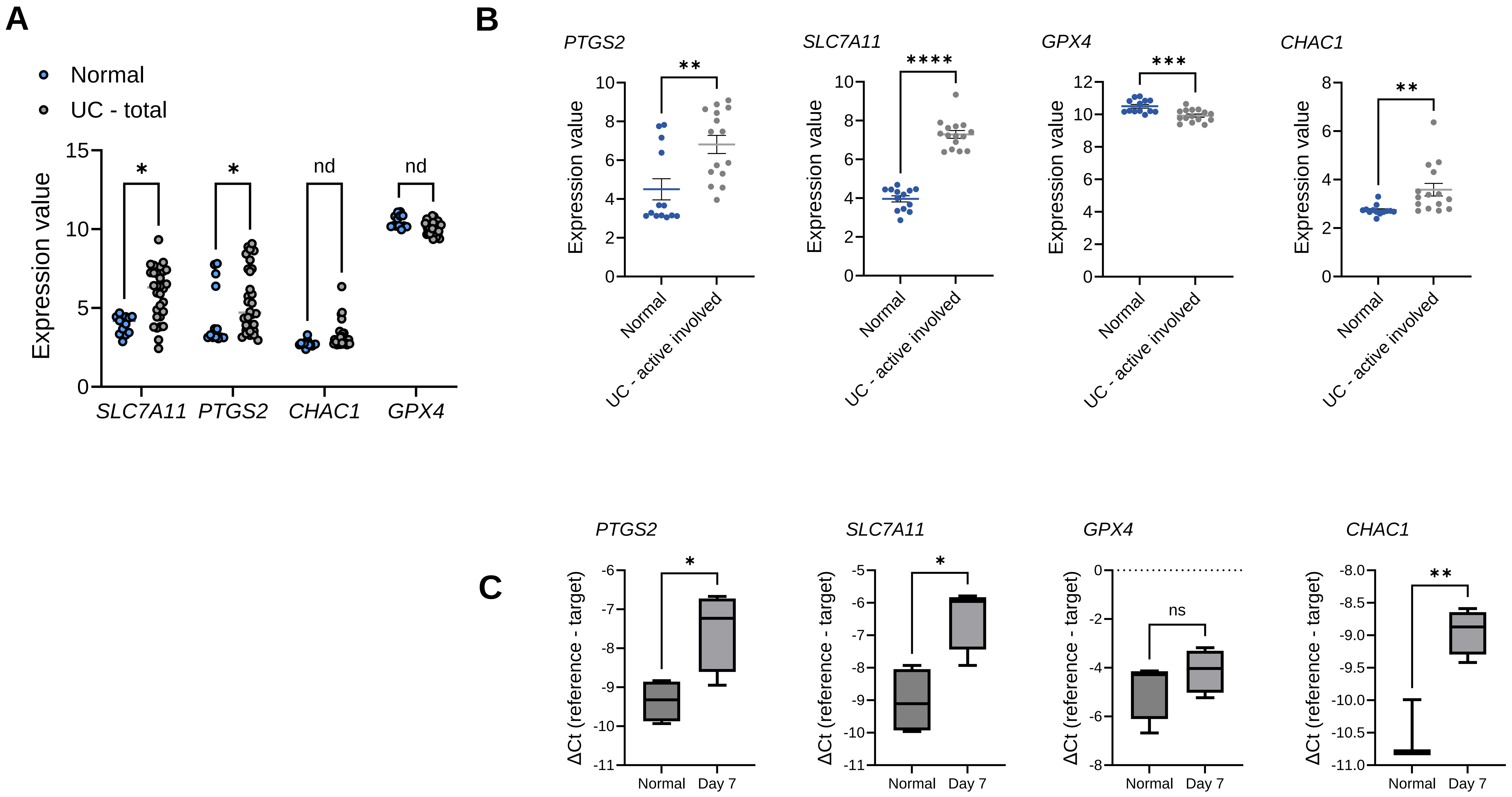


Supplemental Figure 2. (A-B) Human colonic gene expression values derived from GEO data set GSE38713 microarray data. (C) Whole colon mRNA expression by RT-qPCR of tissue from WT mice that were given DSS 2.5% in their drinking water for 5 days, followed by drinking water alone for 2 days after DSS was removed (n = 3-4 mice). Data represent mean ± SEM. *$P < 0.05$, **$P < 0.01$, ***$P < 0.001$, ****$P < 0.0001$ by Student's *t* test and multiple *t* test with correction for multiple comparisons.

### Supplemental Figure 3

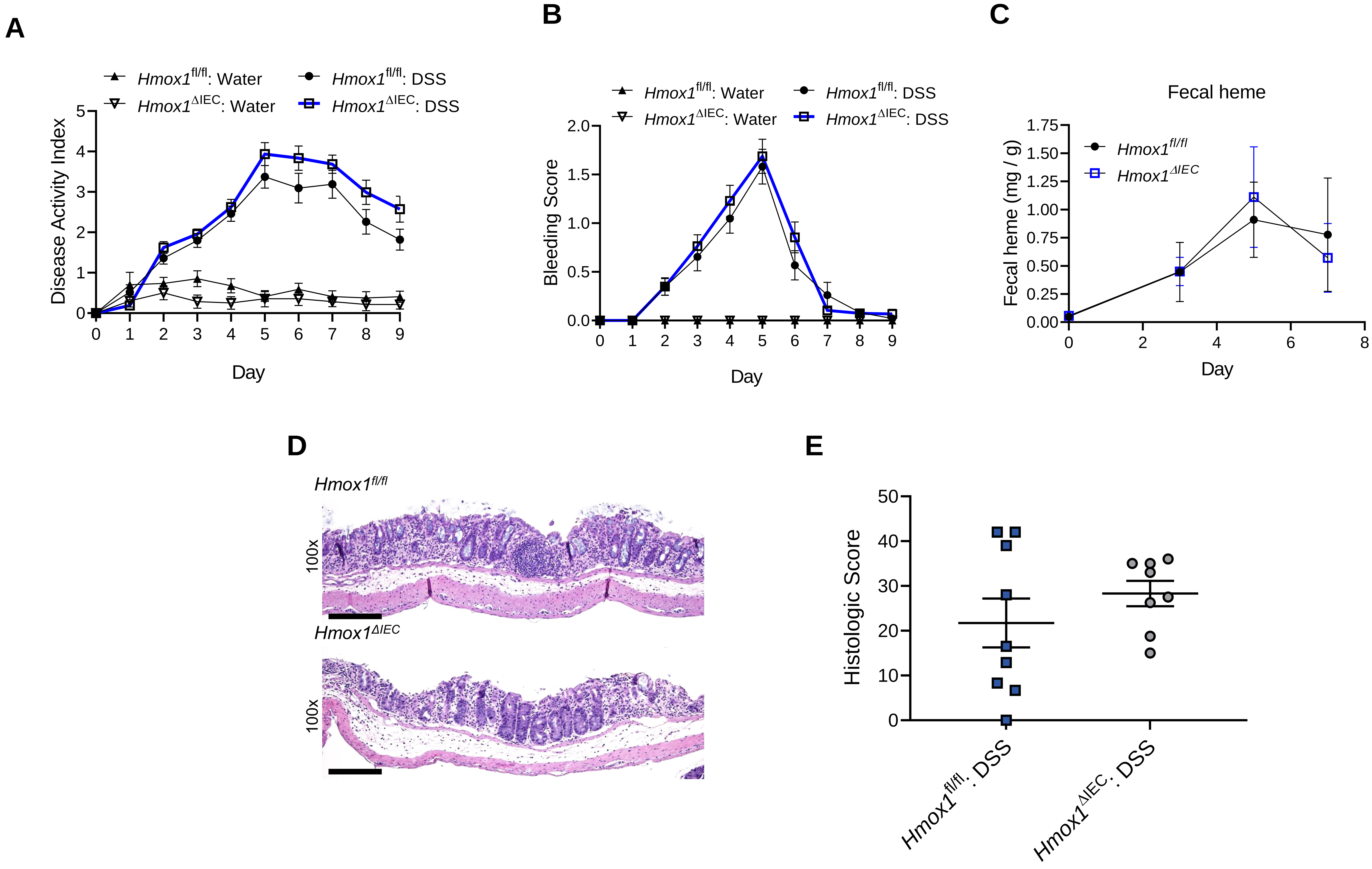


Supplemental Figure 3. (A) DAI scoring of *Hmox1*$^{fl/fl}$ and *Hmox1*$^{\Delta IEC}$ mice (n = 8-11 mice per group from two independent experiments). (B) Fecal bleeding component of DAI scoring of *Hmox1*$^{fl/fl}$ and *Hmox1*$^{\Delta IEC}$ mice. (C) Total fecal heme quantification and DAI bleeding score superimposed from a different cohort of *Hmox1*$^{fl/fl}$ and *Hmox1*$^{\Delta IEC}$ mice undergoing DSS colitis (n = 5-6 mice per group). (D) Representative microscopic images of hematoxylin and eosin (H&E) stained colon tissue at day 7 of a DSS colitis experiment. (E) Histopathological score from colon tissue harvested at day 7 in DSS exposed mice (n = 8-9 mice). Data represent mean ± SEM.

#### Supplemental Figure 4

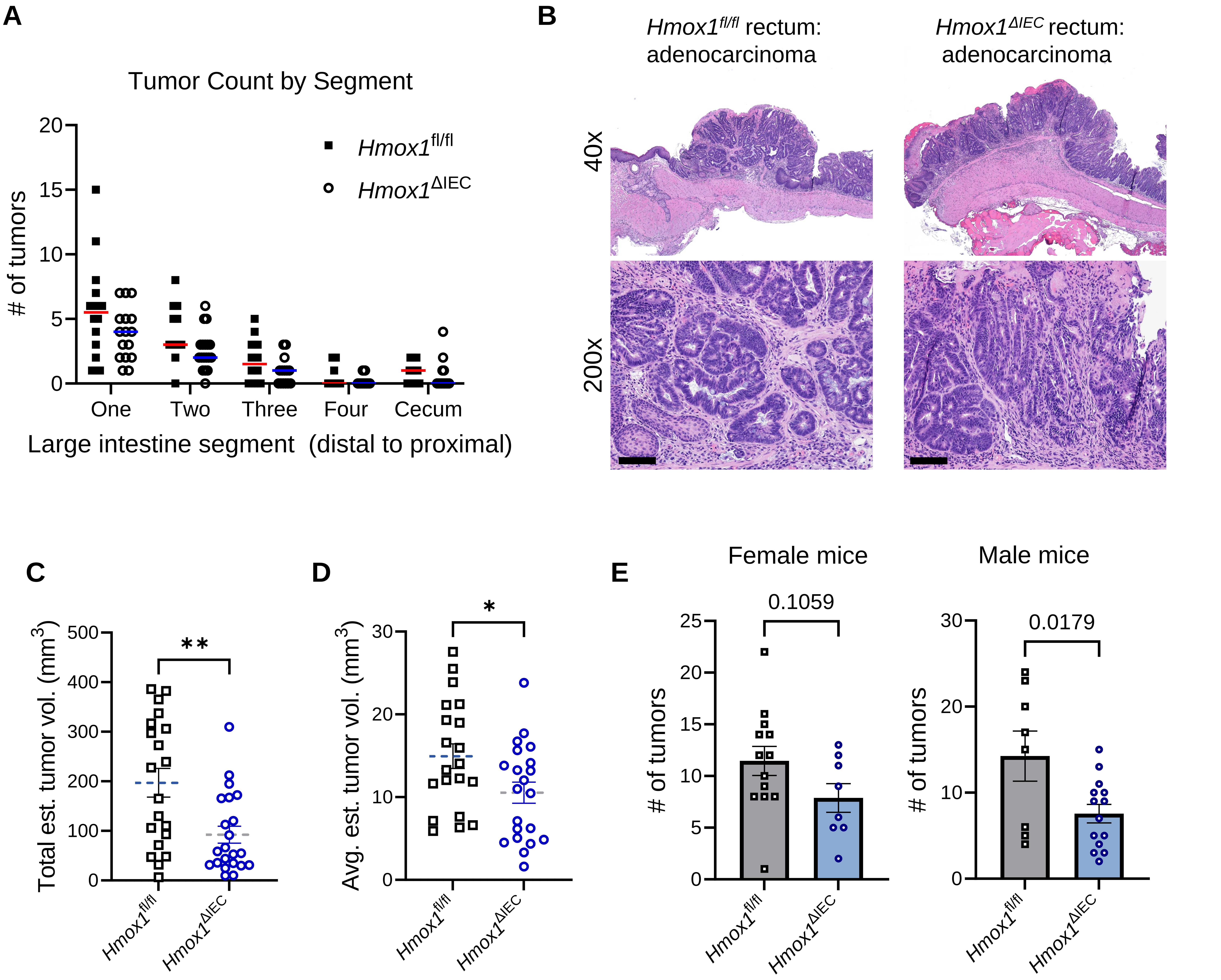


Supplemental Figure 4. AOM-DSS tumor characteristics. (A) Number of tumors per segment starting from the most distal quarter of the colon, proximally to the cecum. (B) Representative H&E stains of rectal adenocarcinomas. (C-D) Tumor volume estimates at day 80 of AOM-DSS experiments obtained by calculating (length * width * width)/2 with (combined results from 3 independent experiments) (E) Total number of tumors when male and female mice are compared separately as a group. Data represent mean ± SEM. *$P$ < 0.05 and **$P$ < 0.01 by unpaired, two-tailed Student's *t* test.

#### Supplemental Figure 5

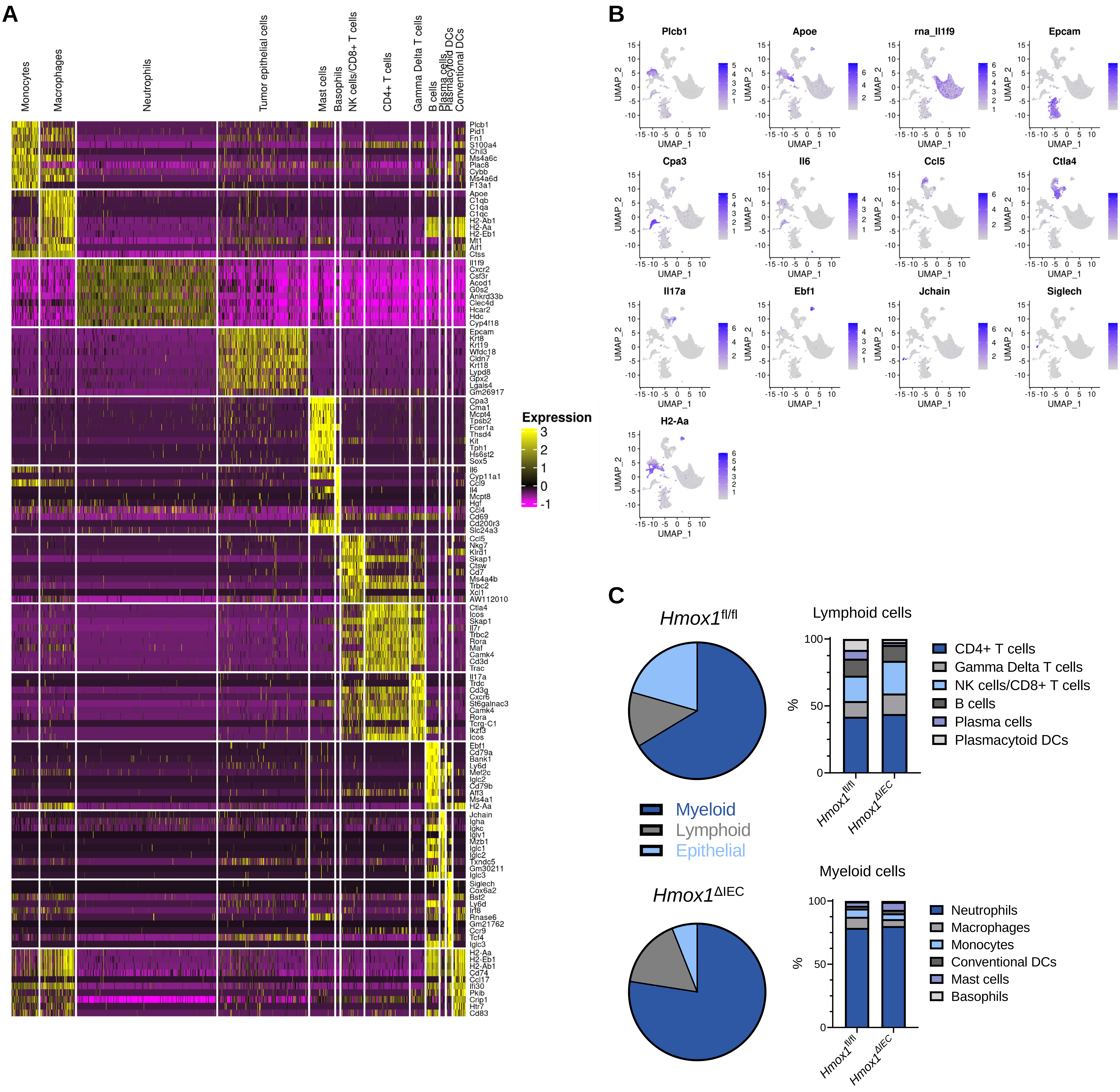


Supplemental Figure 5. (A) Heatmap of top-10 marker genes by cluster, ranked according to $log_2$-fold change multiplied by the difference in percent expression compared to all other cells. (B) UMAP plot highlighting the distribution of the top marker gene expression for each annotated cluster. (C) Pie chart reflecting relative fraction of total analyzed cells composed of myeloid, lymphoid and epithelial populations with bar graphs representing further fractions of the leukocyte subsets.
